# Metabolic demands of the posteromedial default mode network are shaped by dorsal attention and frontoparietal control networks

**DOI:** 10.1101/2022.08.12.503715

**Authors:** GM Godbersen, S Klug, W Wadsak, V Pichler, J Raitanen, A Rieckmann, L Stiernman, L Cocchi, M Breakspear, M Hacker, R Lanzenberger, A Hahn

## Abstract

Although BOLD signal decreases in the default mode network (DMN) are commonly observed during attention-demanding tasks, their neurobiological underpinnings are not fully understood. Previous work has shown decreases but also increases in glucose metabolism that match with or dissociate from these BOLD signal decreases, respectively. To resolve this discrepancy, we analyzed functional PET/MRI data from 50 healthy subjects during the performance of the visuo-spatial processing game Tetris® and combined this with previously published data sets of working memory as well as visual and motor stimulation. Our findings show that the glucose metabolism of the posteromedial DMN is dependent on the metabolic demands of the correspondingly engaged task-positive brain networks. Specifically, the dorsal attention (involved in Tetris®) and frontoparietal networks (engaged during working memory) shape the glucose metabolism of the posteromedial DMN in opposing directions. External attention-demanding tasks lead to a downregulation of the posteromedial DMN with consistent decreases in the BOLD signal and glucose metabolism, whereas working memory is subject to metabolically expensive mechanisms of BOLD signal suppression. We suggest that the former finding is mediated by decreased glutamate signaling, while the latter results from active GABAergic inhibition, regulating the competition between self-generated and task-driven internal demands. The results demonstrate that the DMN relates to cognitive processing in a flexible manner and does not always act as a cohesive task-negative network in isolation.

## Introduction

Large-scale brain networks progressively integrate sensory input to enable complex cognitive processes and behaviors in accordance with internal goals (1). The default mode network (DMN) was originally termed “task-negative” due to decreased blood oxygen level dependent (BOLD) signal (“deactivation”, see Methods for full description) during the performance of external attention-demanding tasks compared to rest (2, 3). Processing of external stimuli in turn increases the BOLD signal in “task-positive” networks including the dorsal attention (DAN) and the frontoparietal network (FPN). While the DAN is mainly engaged when attention to sensory information is required, such as in visuo-spatial reasoning (4), the FPN mediates cognitive control across various task conditions (5) such as the maintenance and manipulation of information also in the absence of external sensory stimuli (6). Because of an anticorrelation in BOLD signals between task-positive and default mode networks at resting-state, it has long been assumed that an antagonism between the DMN and other large-scale networks represents a general characteristic of brain functioning (4). However, the DMN can also be activated during complex cognitive tasks (e.g., memory recollection, abstract self-generated thought) (7, 8). This DMN engagement is thought to facilitate between-network integration, instead of being a segregated network alone (8–10). Moreover, some brain regions such as the posterior cingulate cortex (PCC)/precuneus (part of the posteromedial DMN) have been suggested to play a key role in across-network integration (1). Thus, it has been suggested that task-positive attention/control networks and the DMN can flexibly switch between cooperative and antagonistic patterns to adapt to the task context at hand (1). Despite these advancements in the organization of brain networks, our understanding of the underlying metabolic demands of these context-specific neural processes is limited (11).

Studies using simultaneous fPET/fMRI have shown a strong correspondence between the BOLD signal and glucose metabolism in several task-positive networks and across various tasks requiring different levels of cognitive engagement (12–17). In contrast, a dissociation between BOLD changes (negative) and glucose metabolism (positive) has recently been observed in the DMN during working memory (16), particularly the posteromedial default mode network (pmDMN). In contrast, simple visual and motor tasks elicited a negative metabolic response in this area (18). These findings suggest that distinct underlying metabolic processes support state-specific BOLD signal changes in the DMN (11). However, the consistency and functional specialization of neural and metabolic interactions between default mode and task-positive networks is unclear. Specifically, it is unknown whether the observed dissociation between metabolism and the BOLD signal in the DMN generalizes for complex cognitive tasks, and whether this in turn depends on the brain networks supporting the task performance and their interaction with the DMN.

To address these open questions, we employed functional PET/MR imaging with [_18_F]FDG during rest and performance of a visuo-spatial task requiring a broad range of cognitive functions primarily supported by the dorsal attention network (14, 19). The results were then compared to previously published data from tasks of varying complexity (working memory, eyes opening and finger tapping), primarily involving different brain networks (FPN, visual and motor, respectively). The task response of the posteromedial DMN was the main focus of interest, due to its key role in integrating specialized large-scale brain networks (1, 9). The combination of different imaging modalities and cognitive tasks sheds further light on the interaction across brain networks involved in external attention and control as well as on the corresponding metabolic underpinnings of the BOLD signal.

## Results

Simultaneous fPET/fMRI data were acquired from 50 healthy participants during the performance of the video game Tetris®, a challenging cognitive task requiring rapid visuo-spatial processing and motor coordination (14, 19). From this dataset (DS1) we evaluated the overlap of negative task responses in the cerebral metabolic rate of glucose (CMRGlu) and the BOLD signal specifically in the pmDMN. Next, comparisons to other tasks were drawn, focusing on the corresponding positive task effects across different large-scale functional networks. For this purpose, group-average statistical results of previously published data sets were re-analyzed. These comprised the aforementioned working memory task, specifically the difficult manipulation condition which required active continuation of alphabetic letters (DS2) (16), as well as data from simple eyes open and finger tapping conditions (DS3) (18). After that, the distinct activation patterns across different tasks were used to quantitatively characterize the CMRGlu response of the pmDMN in DS1.. Finally, we investigated the directional influence between the pmDMN and task-positive networks using metabolic connectivity mapping (MCM) (20).

### Regional and task specific effects of CMRGlu and BOLD changes

We first assessed the regional overlap of task-induced changes in CMRGlu and the BOLD signal for the Tetris® paradigm specifically for the DMN. This was directly compared to previous results from the working memory task. The Tetris® task elicited consistent negative responses for both the BOLD signal and CMRGlu in midline core regions of the DMN, such as the medial prefrontal cortex (mPFC) and the PCC/precuneus (DS1, Fig. 1a, all p<0.05 FWE corrected at cluster level, high threshold of p<0.001 uncorrected). This included a cluster in the pmDMN, previously referred to as the ventral PCC (21). In contrast, working memory was associated with a dissociation in the DMN. Here a negative BOLD response was accompanied by increased glucose metabolism in the anterior/dorsal part of the PCC (Fig. 1b; (16)).

**Figure 1:**
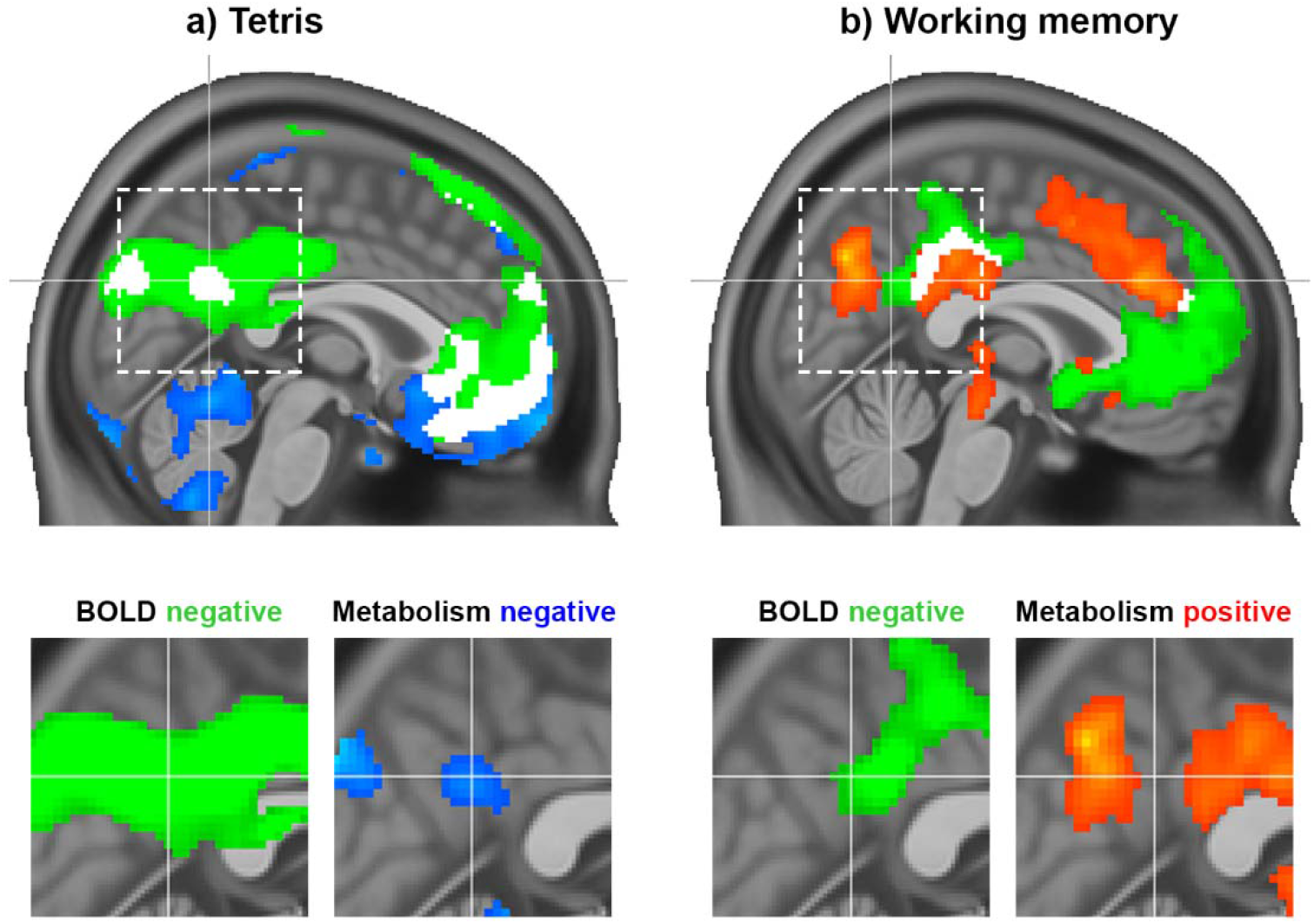
Task responses observed during the Tetris® (DS1) and working memory tasks (DS2) (16). a) The Tetris® task employed in the current work elicited a negative response in the pmDMN for both the BOLD signal (green) and CMRGlu (blue). b) For comparison, previously published results from a working memory manipulation task were also included, which showed a dissociation between BOLD and glucose metabolism in the PCC, i.e., negative BOLD response (green) vs. increased metabolism (red). White clusters represent the intersection of significant CMRGlu and BOLD signal changes, irrespective of direction. The dashed rectangle indicates the zoomed section of the PCC. All modalities are corrected for multiple comparisons (p<0.05). Crosshair is at −1/-56/30 mm MNI-space.

We then compared task-induced changes across all task paradigms, evaluating relationships between task-positive and task-negative effects. Using a common cortical parcellation scheme of seven functional networks (22), this analysis revealed distinct spatial patterns for the different tasks (Fig 2). Tetris® elicited positive changes for BOLD and CMRGlu predominantly in the visual network (VIN) and DAN, while working memory mostly involved the FPN. The greatest difference in positive task effects between Tetris® and working memory was therefore observed for VIN, DAN and FPN (white bars in Fig 2c), which were selected for further evaluation. Overlapping negative responses for both imaging modalities occurred in DMN3 (“core”) and DMN4 (“ventral”) for Tetris®, but only in DMN4 for working memory (Fig 2f). Simple visual stimulation (eyes open vs eyes closed) and right finger tapping elicited increased CMRGlu in VIN and SMN (somato-motor), respectively, and a negative response mostly in DMN3 (Suppl Fig 1). These results were reproduced when computing the overlap between imaging modalities by statistical conjunction analysis in SPM12 (Suppl Fig 1). Thus, the above results were obtained on the basis of common task-specific changes between the BOLD signal and CMRGlu. For completeness, we also evaluated the overall regional agreement between the two imaging modalities, confirming that activations (Dice coefficient Tetris®=0.57 and working memory=0.35) showed higher overlap than deactivations (Tetris®=0.16 and working memory=0.06) (16).

**Figure 2:**
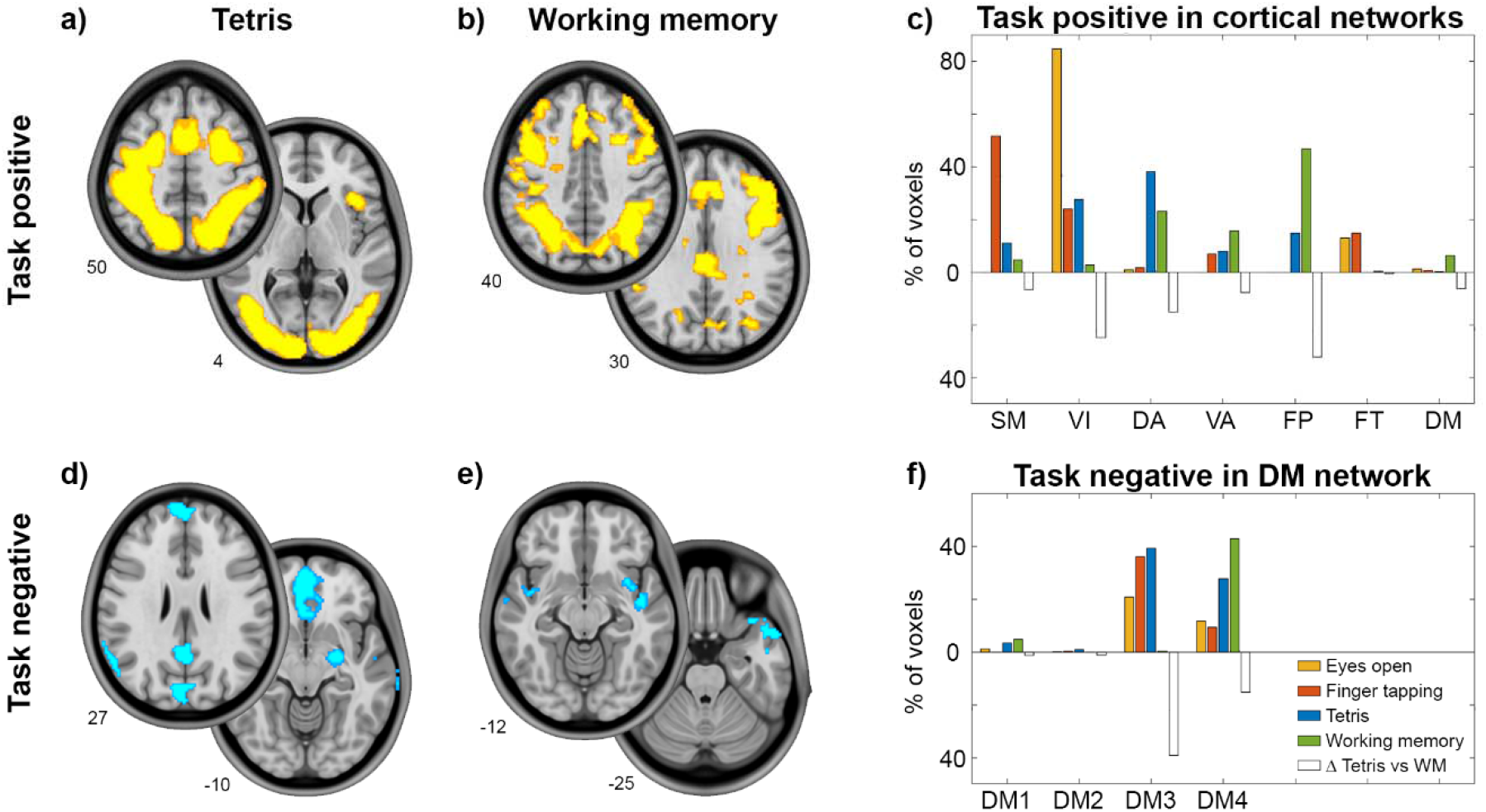
Detailed response for the cognitive tasks. Task effects represent the overlap computed as the intersection between BOLD signal changes and glucose metabolism (all p<0.05 corrected). Slices show major clusters of positive and negative responses for the two tasks and numbers indicate the z-axis in mm MNI space. Bar graphs show the percentage of voxels with a positive task response for each of the 7 cortical networks (22). a-c) Differences in positive task responses between Tetris® (DS1) and working memory (DS2) (16) were most pronounced in visual, dorsal attention and fronto-parietal networks, as indicated by open bars (absolute difference between Tetris and working memory). For completeness, previous CMRGlu data obtained while opening the eyes (orange) and right finger tapping (red) was also included (DS3) (18). These elicited the main task-positive response in visual and somato-motor networks, respectively. d-f) Negative task responses are shown for DMN subparts as given by the 17-network parcellation (22), with DMN3 and DMN4 covering mostly core (PCC, mPFC, angular) and ventral areas (temporal, lateral OFC, superior frontal), respectively. The negative response was strongest in DMN3 for Tetris®, visual and motor tasks, but in DMN4 for the working memory task. The number of voxels per network was normalized by the total number of activated (c) or deactivated (f) voxels across both imaging modalities. Thus each task sums up to 100% across all cortical networks.

### Task positive CMRGlu shapes pmDMN response

We next characterized the negative CMRGlu response in the pmDMN observed in DS1. Individual CMRGlu values were extracted for this region and from the networks that showed the greatest difference between Tetris® and the working memory tasks (VIN, DAN and FPN). These were then entered in a linear multiple regression analysis.

The negative CMRGlu task response in the pmDMN during Tetris® significantly covaried with the positive CMRGlu response in the other networks (Fig 3a, F=4.84, p=0.005). More specifically, CMRGlu of the pmDMN was associated with that of FPN (b=0.75, p=0.006) and DAN (b=-0.68, p=0.010), but not VIN (b=0.30, p>0.19). Notably, the influence of DAN and FPN was in the opposite direction (as given by the opposite sign of the parameter estimates). That is, across individuals, the combination of low glucose metabolism in FPN and high metabolism in DAN was associated with a negative CMRGlu task response in the pmDMN. Similar results were obtained when defining the overlap across imaging modalities as a formal statistical conjunction (whole model: F=3.32, p=0.028; FPN: b=0.39, p=0.018, DAN: b=-0.55, p=0.011, VIN: b=0.17, p=0.33). The association was even stronger when using the atlas-based network definition (22), i.e., without incorporating prior knowledge of positive task responses (whole model: F=6.65, p=0.0008, FPN: b=1.04, p=0.002, DAN: b=-0.98, p=0.001, VIN: b=0.69, p=0.030). The influence of the visual network also reached significance when using the atlas definition, in line with our initial work showing CMRGlu decreases in the pmDMN when comparing eyes opened vs. eyes closed (18).

**Figure 3:**
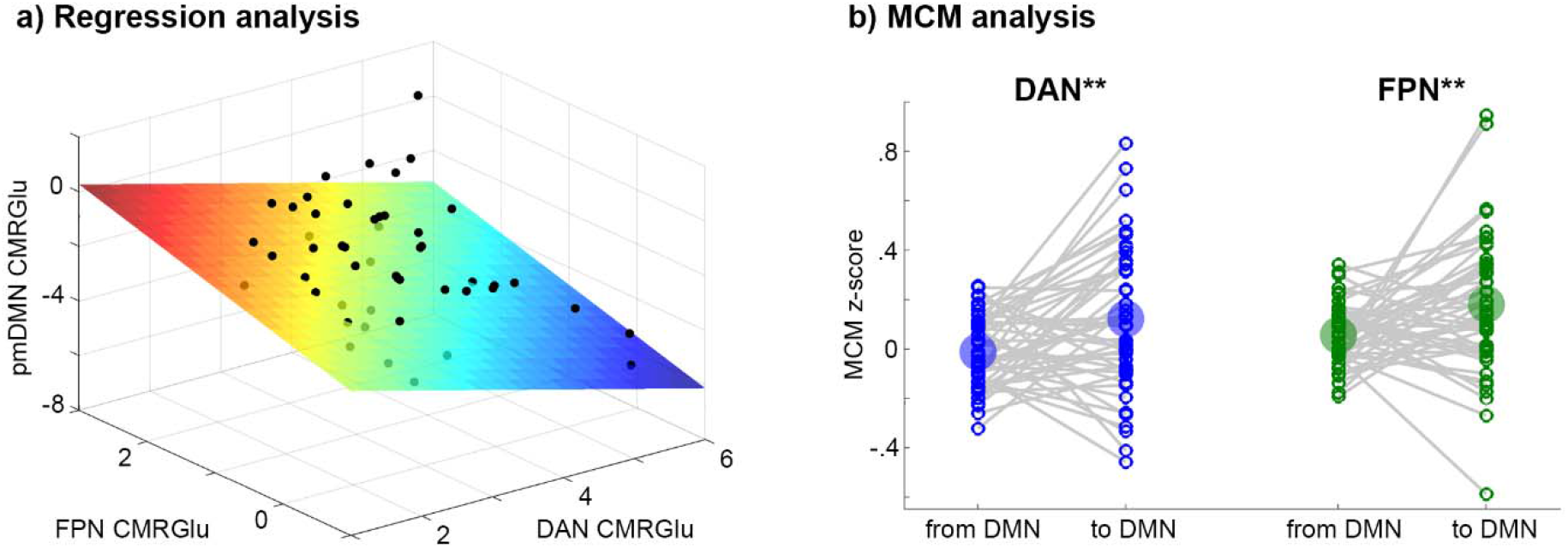
Relationship of CMRGlu response between networks. a) Visualization of regression analysis results for the Tetris® task (DS1, F=4.84, p=0.005). Positive CRMGlu task responses of the FPN (p=0.006) and the DAN (p=0.010) both explained the negative CMRGlu response in the pmDMN across subjects. Here, FPN and DAN exerted an inverse association with pmDMN, where low glucose metabolism in FPN and high metabolism in DAN yield a negative CMRGlu response in the pmDMN. Units on all axes are µmol/110g/min. b) MCM analysis for the Tetris task (DS1) combining the association of functional connectivity and CMRGlu for causal inference on directionality. The influence from DAN and FPN to DMN was significantly stronger than vice versa (both **p<0.01). The same regions were used for regression and MCM analyses: For FPN and DAN CMRGlu was extracted from voxels showing a significant overlap of activations between imaging modalities but no overlap between Tetris® and WM tasks.

Finally, MCM was employed to assess the putative direction of influence between task-positive networks and the DMN. The technique is based on previous evidence showing that most energy demands emerge postsynaptically (23–25), which is used to identify the target region of a connection. Applied to imaging parameters, this is reflected in a correlation of spatial patterns between functional connectivity and the underlying CMRGlu, thereby enabling causal inferences (see Methods). In line with the regression analysis, MCM indicated that during task performance the direction of influence is from DAN (MCM z-score=0.12 vs −0.01, p=0.005, Fig 3b) and FPN (MCM=0.18 vs 0.05, p=0.006) to the pmDMN.

## Discussion

Using simultaneous PET/MR imaging, we find that the metabolic demands linked to pmDMN BOLD deactivations depend on the actual task at hand (external vs. internal processing) and the correspondingly activated functional networks (visual/motor and attention vs. control). Specifically, we showed that high task-induced metabolism in the DAN but low metabolism in the FPN led to a negative CMRGlu response in the pmDMN. These findings resolve the discrepancy between (non)congruent energy demands and BOLD signal changes across different cognitive tasks, emphasizing the distinct metabolic underpinnings of BOLD signal deactivations during cognitive processing.

### Spatially distinct metabolic response within the DMN

The tasks investigated in this work resulted in distinct metabolic responses in the pmDMN (Fig 1-2). This included the PCC, a spatially and functionally heterogeneous region (26) with ventral and dorsal subcomponents (vPCC/dPCC) (27). The PCC is a central hub, connecting brain networks involved in complex behavior (28). Specifically, the PCC is thought to act as an interface between distributed functional networks by echoing their activity (21) and tuning the balance between the internal and external focus of attention (29). Despite a consistent negative BOLD response during external attention-demanding tasks, the dPCC and vPCC are differently integrated into the DMN and distinct large-scale networks underpinning task performance (21, 30). That is, these two DMN regions are thought to be flexibly recruited to support various cognitive functions (1).

The vPCC is engaged in tasks with an internal focus of attention (29), such as autobiographical memory recollection (31), and when demands for externally directed attention are low (21). Our findings add to this knowledge by suggesting that vPCC BOLD deactivation represents a downregulation of internal processing in favor of a focus on the external attention-demanding task e.g., as supported by DAN, VIN and SMN (21). Thus, a negative BOLD response paralleled by a decreased CMRGlu, as observed for Tetris® as well as simple visual and motor stimulation, indicates a reduction of overall metabolism, in line with the original understanding of DMN task-negativity (2, 4, 32).

In contrast, the dPCC is involved in externally directed attention and plays an opposite role to the vPCC (21). With increasing working memory load, the dorsal subregion exhibits stronger integration with the DMN and more pronounced BOLD signal anticorrelation with task-control networks, which is also reflected in behavioral performance (21). This could potentially lead to a competition between self-generated and task-induced internal demands. Thus, vPCC BOLD deactivation during working memory seems to reflect a metabolically expensive process that suppresses self-generated thoughts to enhance task focus, particularly when internal task demands are high (7, 16, 21, 33).

The associations of these metabolic demands between the DMN and task-positive networks is also reflected in their distance along a connectivity gradient, which is hierarchically organized from unimodal sensory/motor to complex associative functions and the DMN being at the end of the processing stream (8, 34). A corresponding decrease in pmDMN glucose metabolism was observed for tasks that activate unimodal networks and the DAN, but not for the FPN. The inverse influence of attention and control networks on the pmDMN may therefore suggest that connectivity gradients are supported by the underlying energy metabolism.

### Underlying neurophysiological effects

The distinct relationships between BOLD and CMRGlu signals that emerge during specific tasks highlight the different physiological processes contributing to neuronal activation of cognitive processing (11). Although not directly measured in the current work, we aim to relate our findings to neurotransmitter actions that could potentially explain the measured imaging parameters. Previous studies suggest that BOLD decreases in DMN are associated with excitatory-inhibitory neurotransmitter balance (35), with a lower glutamate/GABA ratio leading to greater PCC deactivation (36). Being the major excitatory neurotransmitter, glutamate increases the BOLD signal upon release and thus prevents negative BOLD responses in the PCC (35). Moreover, glutamate signaling leads to a proportional glucose consumption (37, 38), which provides energy for reversal of ion fluxes (23). Thus, decreases in glutamate signaling in the pmDMN could explain decreased BOLD and CMRGlu signals during the Tetris® task.

Conversely, the major inhibitory neurotransmitter GABA supports a negative BOLD response in the PCC (35). Although the influence of GABA on CBF and thus the BOLD signal is not straightforward (39), BOLD decreases in the PCC have been specifically related to increased GABA signaling during a working memory task (40) as well as upon pharmacological stimulation with diazepam, a positive allosteric modulator of GABA (41). This GABAergic inhibition may require active glucose consumption to restore membrane potentials (although to a lesser extent than glutamate signaling (23)). Indeed, specific GABAergic neurons (i.e., parvalbumin-containing interneurons) are particularly metabolically demanding (42) and involvement of this cell type would be in line with the suppression of characteristic gamma oscillations (43).

In this context, the observed dissociation between BOLD changes and CMRGlu during working memory could result from metabolically demanding GABAergic suppression. At the same time, the correspondence of both imaging parameters during Tetris® relates to a reduction of glutamatergic input. We, therefore, speculate that the effect of the DAN on the pmDMN may be mediated via glutamate, while the effect of the FPN relates to GABA, resulting in congruent and opposing changes in BOLD and CMRGlu signals, respectively. Future work may test this hypothesis, using the pharmacological alteration of GABAergic and glutamatergic signaling or optogenetic approaches modulating GABAergic interneuron activity.

### Limitations, outlook and conclusions

To summarize, our work provides novel insights into the metabolic underpinnings of negative BOLD responses in the pmDMN, showing that regionally specific effects depend on the functional networks involved in task execution. Acknowledging the underlying energy demands and neurotransmitter actions underpinning neural activity is necessary to understand PCC function, including how this essential DMN region dynamically interacts with macroscale networks as a function of changing behavioral demands (44, 45). While our work provides valuable information to address this knowledge gap, several caveats are worth noting: First, we did not assess the different tasks in the same cohorts, but pooled different studies. However, only effects corrected at the group level were used as obtained from commonly employed sample sizes, which should provide representative findings. Second, Tetris® and WM data were acquired with different task designs, i.e. continuous task performance versus hierarchical embedding of short task blocks for BOLD into longer PET acquisition, respectively. As the latter may not clearly differentiate between start-cue and task activation, this may limit transferability. Therefore, future studies investigating these effects should address this limitation, ideally studying the different tasks in the same cohort, with a comparable task design. Third, our results were obtained using rest with cross-hair fixation as the baseline condition. Since task-specific effects in the BOLD signal and CMRGlu are relative to this baseline condition, these would likely change if using an active control condition as baseline. Fourth, additional contextual load input to the pmDMN, (e.g. task-relevant emotional content) may be another important factor affecting its activation (46). Future studies may therefore include additional cognitive and emotional domains. Of particular interest would be the investigation of introspective tasks such as autobiographical memory as these typically induce a positive BOLD response in the PCC, while the coupling with the CMRGlu response is unknown. Although the DMN, and in particular the PCC, has been implicated in numerous brain disorders (33), our data suggests that this could be mediated by other interacting cortical networks. Assessing the differential influence of attention and control networks on the pmDMN may therefore represent an interesting approach to improve our understanding of network dysfunction in different patient populations.

## Methods

Throughout this manuscript we refer to deactivation/decrease/negative response and likewise to activation/increase/positive response as relative changes compared to the baseline condition. That is, changes in the BOLD signal and glucose metabolism (here as obtained from general linear model analyses) emerge from a negative or positive contrast sign with respect to the baseline. For glucose metabolism, these changes are absolutely quantified in µmol/100g/min.

In this work the baseline condition for all tasks was a resting-state defined as looking at a crosshair without focusing on anything in particular, except for the eyes open condition which used eyes closed as baseline (please see discussion for implications when other control conditions are employed).

### Data sets

The primary dataset used in this work (DS1) consists of simultaneously acquired fPET/fMRI data from n=50 healthy subjects during the performance of a challenging visuo-motor task (i.e. the video game Tetris®). Unless stated otherwise, all data, methods and results refer to DS1, including data of individual participants and all statistics across subjects. A detailed description of the study design, cognitive task, fPET/fMRI measurements and first-level analysis of DS1 is given in our previous work (14) and also below.

For direct comparison with previous work, two further data sets were used in this study. These only include group-averaged results with contrasts and statistical maps as published previously. Dataset 2 (DS2) consists of fPET/fMRI data from n=23 healthy subjects (mean age ± sd = 25.2 ± 4.0 years, 13 female) acquired during the performance of a working memory manipulation task, which required active transformation of stimuli in working memory (16). Group-level maps of significant increases and decreases in [_18_F]FDG glucose metabolism and the BOLD signal during task execution as compared to baseline were used (p<0.05 TFCE corrected). Dataset 3 (DS3) comprises data on glucose metabolism obtained from [_18_F]FDG fPET imaging (18). We included group-average statistical maps from n=18 healthy subjects (24.2 ± 4.3 years, 8 females), who performed cognitively simple tasks of eyes open vs. eyes closed and during right finger tapping vs. rest (p<0.05 FWE corrected cluster-level after p<0.001 uncorrected voxel level).

For all datasets and tasks, the baseline condition was defined as looking at a crosshair at resting-state, except for the eyes open condition of DS3 which used eyes closed as the baseline.

### Experimental design

All participants of DS1 underwent one PET/MRI scan while performing a challenging visuo-spatial cognitive task. Data acquisition started with structural imaging (8 min). This was followed by 52 min fPET, which comprised an 8 min baseline at rest and then four periods of continuous task performance (6 min each, two easy and two hard conditions, randomized) with periods of rest after each task block (5 min). Simultaneously with fPET, BOLD fMRI was acquired during the continuous task execution (6 min each), which was used for the calculation of metabolic connectivity mapping. Finally and immediately after fPET, another BOLD fMRI sequence was obtained in a conventional block design with the same task (12 task blocks, 30 s each, four easy, four hard and four control blocks, 10 s baseline between task blocks, 8.17 min in total). This acquisition was used for the computation of BOLD-derived neuronal activation. Further acquisitions (diffusion weighted imaging, BOLD imaging at rest, arterial spin labelling) were not used in the current work. During all periods of rest, participants were instructed to look at a crosshair, relax and not to focus on anything in particular.

### Cognitive task

An adapted version of the video game Tetris® (https://github.com/jakesgordon/javascript-tetris, MIT license) was implemented in electron 1.3.14. The aim is to build complete horizontal lines by rotation and alignment of bricks, which descend from the top of the screen. The task included two levels of difficulty, which differed regarding the speed of the descending bricks (easy/hard: 1/3 lines per sec) and the number of incomplete lines built at the bottom (easy/hard: 2/6 lines out of 20). The control condition of the BOLD acquisition was not used in this work. Right before the start of the PET/MRI scan, participants familiarized themselves with the control buttons by 30 s training of each task condition. The employed task represents a cognitively challenging paradigm, which requires a high level of attention, rapid visuo-spatial motor coordination, mental rotation, spatial planning and problem solving.

### Participants

Fifty-three healthy participants were initially recruited for DS1 and 50 were included in the current analysis (mean age ± sd = 23.3 ± 3.4 years, 23 female). Reasons for drop out were failure of arterial blood sampling (n=1) and technical issues during the scan (n=2). In part these subjects also participated in previous studies (14, 19, 47). All subjects completed an initial screening to ensure general health through a routine medical examination (blood tests, electrocardiography, neurological testing, structural clinical interview for DSM-IV). Female participants also underwent a urine pregnancy test at the screening visit and before the PET/MRI examination. Exclusion criteria were current or previous somatic, neurological or psychiatric disorders (12 months), substance abuse or psychopharmacological medication (6 months), current pregnancy or breast feeding, contraindications for MRI scanning, previous study-related radiation exposure (10 years) and previous experience with the video game Tetris® (3 years). All participants provided written informed consent after a detailed explanation of the study protocol, they were insured and reimbursed for participation. The study was approved by the Ethics Committee of the Medical University of Vienna (ethics number 1479/2015) and procedures were carried out according to the Declaration of Helsinki. The study was pre-registered at ClinicalTrials.gov (NCT03485066).

### PET/MRI data acquisition

Participants had to fast for at least 5.5 hours before the start of the PET/MRI scan, except for unsweetened water. The radiotracer [_18_F]FDG was applied in a bolus + infusion protocol (510 kBq/kg/frame for 1 min, 40 kBq/kg/frame for 51 min) using a perfusion pump (Syramed µSP6000, Arcomed, Regensdorf, Switzerland), which was kept in an MRI-shield (UniQUE, Arcomed).

MRI acquisitions included a T1-weighted structural scan (MPRAGE sequence, TE/TR = 4.21/2200 ms, TI = 900 ms, flip angle = 9°, matrix size = 240 × 256, 160 slices, voxel size = 1 × 1 × 1 mm + 0.1 mm gap, 7.72 min) and BOLD fMRI (EPI sequence, TE/TR = 30/2000 ms, flip angle = 90°, matrix size = 80 × 80, 34 slices, voxel size = 2.5 × 2.5 × 2.5 mm + 0.825 mm gap, 6 min for functional connectivity and 8.17 min for neuronal activation in the block design).

### Blood sampling

Before the PET/MRI scan blood glucose levels were assessed as triplicate (Glu_plasma_). During the PET/MRI acquisitions manual arterial blood samples were drawn at 3, 4, 5, 14, 25, 36 and 47 min after the start of the radiotracer administration (15). From these samples whole-blood and plasma activity were measured in a gamma counter (Wizard_2_, Perkin Elmer). The arterial input function was obtained by linear interpolation of the manual samples to match PET frames and multiplication with the average plasma-to-whole-blood ratio.

### Cerebral metabolic rate of glucose metabolism (CMRGlu)

PET images were reconstructed and processed as described previously (15). Briefly, PET list mode data were corrected for attenuation with a database approach (48) and reconstructed to 30 s frames (matrix size = 344 × 344, 127 slices). Preprocessing was carried out in SPM12 (https://www.fil.ion.ucl.ac.uk/spm/) and comprised motion correction (quality = 1, register to mean), spatial normalization to MNI space using the T1-weighted structural image and spatial smoothing with an 8 mm Gaussian kernel. Non-gray matter voxels were masked out and a low pass filter was applied (cutoff frequency = 3 min). Identification of task-specific effects was done with the general linear model. Four regressors were included to characterize the baseline, the two task conditions (easy and hard as a linear function with slope = 1 kBq/frame) and head motion (the first principal component of the 6 motion parameters). The baseline regressor was given by the average time course of all gray matter voxels, excluding those activated during the hard task condition of the individual BOLD block design (p< 0.05 FWE corrected voxel level). This approach provides the best model fits (15) without negatively affecting task-specific changes in glucose metabolism (14, 15) or test-retest reliability (47). Quantification was carried out with the Patlak plot and the influx constant K_i_ was converted to CMRGlu as

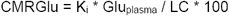

with LC being the lumped constant = 0.89. The resulting maps represented individual task-specific changes in CMRGlu, which were used for group-level analyses and MCM.

### Blood oxygen level dependent (BOLD) signal changes

BOLD imaging data of the block design were processed with SPM12 as described previously (15). In short, data were corrected for slice timing effects (reference = middle slice) and head motion (quality = 1, register to mean), normalized to MNI-space via the T1-weighted image and smoothed with an 8 mm Gaussian kernel. First-level analysis was carried out to assess individual estimates of the BOLD task response. Here, regressors were included for the two task conditions (easy, hard) and the control condition as well as nuisance regressors for head motion, white matter and CSF signals. The contrast of interest was chosen as the hard task level vs. baseline to facilitate comparison to previous work (16).

### Metabolic connectivity mapping (MCM)

MCM was calculated during the performance of the hard task between regions of interest as published previously (14). The approach combines patterns of functional connectivity (FC) and CMRGlu and enables to estimate directional connectivity between brain regions (20). MCM relies on physiological characteristics of energy demands, which mostly emerge postsynaptically (23, 24) and thereby allow to identify the target region of a connection. Briefly, computing the FC between region A and region B yields a distinct FC voxel-wise pattern in region B. If the influence of region A on B is causal, this will result in a corresponding CMRGlu pattern in region B because of the coupling between the BOLD signal and the underlying glucose metabolism (49, 50). FC was computed from continuously acquired BOLD imaging data during the performance of the hard task condition. Preprocessing was done as described above for the BOLD block design. After spatial smoothing, motion scrubbing was performed by removing frames with a displacement > 0.5 mm (plus one frame back and two forward). To remove potentially confounding signals linear regression was used (including motion parameters, white matter and cerebrospinal fluid signals), followed by band-pass filtering (0.01 < f <0.15 Hz (51)). FC was then calculated as the temporal correlation between regions A and B. MCM was computed as the spatial correlation between voxel-wise patterns of FC and CMRGlu in region B (both correlations were z-transformed). ROIs were defined as for the regression analysis below, i.e., as voxels of the DAN and FPN which were specific for each task as well as voxels of the pmDMN identified with DS1. One subject was excluded from MCM analysis because of movement during the BOLD acquisition of the hard task condition.

### Statistical analysis

For DS1, task responses in CMRGlu and BOLD were evaluated during the Tetris® task with separate one sample t-tests for each imaging modality (p<0.05 FWE corrected cluster-level after p<0.001 uncorrected voxel level). Group maps with significant changes were then binarized and the overlap between CMRGlu and BOLD responses was computed as their intersection. First, this was done for negative task effects to assess if DMN deactivations are present in both modalities.

Next, we tested the hypothesis that negative responses in the DMN are mirrored by the corresponding positive task changes that are specific to other cortical networks. To provide a comprehensive overview of changes across different tasks we also included results from a working memory task (DS2) (16) as well as eyes opened vs. eyes closed conditions and right finger tapping (DS3) (18). For the Tetris® and working memory tasks, overlapping increases in CMRGlu and BOLD signal changes were computed in the same way as above, i.e. by computing the intersection of group-level significant positive task effects across modalities for each task separately. For the visual and motor paradigms, only CMRGlu data was available. The different positive task responses were first compared qualitatively by computing the percentage of activated voxels for each task and 7 cortical networks (22). Negative task responses were calculated at a finer level of detail with the DMN subdivisions obtained from the 17 network parcellation (22).

After that, these network-specific differences in positive responses between the Tetris® and working memory tasks (i.e. DS1 vs DS2) were then used to quantitatively explain the CMRGlu decreases in the pmDMN observed in the current dataset. Individual CMRGlu values of DS1 were extracted for visual, dorsal attention and fronto-parietal networks, as these networks showed greatest differences in activation (see results). Here, only voxels were used which showed an overlap of significant task changes (i.e., the intersection between CMRGlu and BOLD signal) and which were specific for each task (i.e., voxels that did not overlap between the two tasks). Voxels of the pmDMN were defined by the overlapping negative task response between CMRGlu and BOLD changes of DS1. The extracted values were then entered into a regression analysis to characterize the CMRGlu response in the pmDMN. To assess the robustness of our results and allow for generalization, two control analyses were performed. First, the overlap of task responses across imaging modalities at the group-level was also computed by a formal conjunction analysis in SPM12 for the Tetris® and working memory tasks. That is, for each task the individual maps of CMRGlu and BOLD response were z-scored and entered in a one-way ANOVA with each modality representing a “group”. The separate contrasts for each of the modalities were then combined in a conjunction (p<0.05 FWE corrected cluster level following p<0.001 uncorrected voxel level). Second, the regression was also calculated when extracting CMRGlu values from the entire visual, dorsal attention and fronto-parietal networks (22), i.e. independent of regionally specific prior knowledge on positive task responses.

Finally, we assessed the directionality of the association between DMN and task-positive networks in DS1. Individual MCM values were entered in a paired t-test to assess differences between directions, e.g., DAN->DMN vs. DMN->DAN.

## Acknowledgements

We thank the graduated team members and the diploma students of the Neuroimaging Lab (NIL, head: R. Lanzenberger) as well as the clinical colleagues from the Department of Psychiatry and Psychotherapy for clinical and/or administrative support. In detail, we would like to thank S. Kasper, K. Papageorgiou, P. Michenthaler, T. Vanicek, A. Basaran, M. Hienert, L. Silberbauer, J. Unterholzner and G. Gryglewski medical support, L. Rischka and M. B. Reed for analysis support, V. Ritter, K. Einenkel and E. Sittenberger for subject recruitment and A. Jelicic for partly implementation of the task. We are further grateful to J. Völkle and A. Pomberger for radioligand synthesis. The scientific project was performed with the support of the Medical Imaging Cluster of the Medical University of Vienna.

This research was funded in whole, or in part, by the Austrian Science Fund (FWF) KLI 610, PI: A. Hahn. For the purpose of open access, the author has applied a CC BY public copyright license to any Author Accepted Manuscript version arising from this submission. S. Klug is supported by the MDPhD Excellence Program of the Medical University of Vienna.

A.R. and L.S. are supported by the European Research Council under the European Union’s Horizon 2020 research and innovation program (ERC-STG-716065 to A.R). LC is supported by the Australian NHMRC (GN2001283).

## Author contributions

Study design: A.H., R.L., M.H.

Data acquisition: G.M.G., A.H., S.K., V.P., W.W., J.R.

Data analysis: A.H., L.S.

Reagants / Methods: A.H., L.S., A.R., M.B., L.C.

Manuscript preparation: G.M.G., A.H., L.C., M.B., L.S., A.R.

All authors discussed the implications of the findings and approved the final version of the manuscript.

## Competing interests

R. Lanzenberger received travel grants and/or conference speaker honoraria within the last three years from Bruker BioSpin MR and Heel, and has served as a consultant for Ono Pharmaceutical. He received investigator-initiated research funding from Siemens Healthcare regarding clinical research using PET/MRI. He is a shareholder of the start-up company BM Health GmbH since 2019. M. Hacker received consulting fees and/or honoraria from Bayer Healthcare BMS, Eli Lilly, EZAG, GE Healthcare, Ipsen, ITM, Janssen, Roche, and Siemens Healthineers. W. Wadsak declares to having received speaker honoraria from the GE Healthcare and research grants from Ipsen Pharma, Eckert-Ziegler AG, Scintomics, and ITG; and working as a part time employee of CBmed Ltd. (Center for Biomarker Research in Medicine, Graz, Austria). All other authors report no competing interests in relation to this study.

## Data availability

Raw data will not be publicly available due to reasons of data protection. Processed data can be obtained from the corresponding author on request with a data-sharing agreement.

## Code availability

Custom code can be obtained from the corresponding author on request.

