## Supplementary material for "Metabolic demands of the posteromedial default mode network are shaped by dorsal attention and frontoparietal control networks": Suppl Fig 1

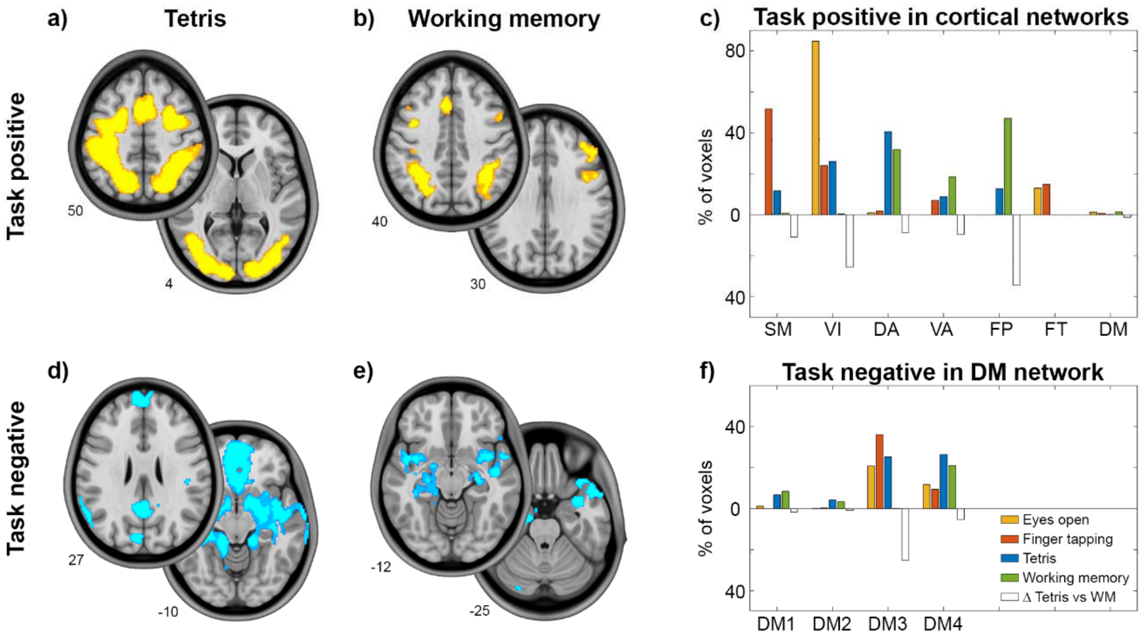


***Supplementary Figure S1: Task response computed by statistical conjunction analysis.*** For direct comparison with figure 2, the overlap between CMRGlu and BOLD signal changes was computed by conjunction analysis in SPM12 (all $p<0.05$ FWE corrected cluster-level after $p<0.001$ uncorrected voxel level). Task-specific effects were similar to the intersection between imaging modalities: The Tetris® paradigm showed increases in imaging parameters, mostly for VIN and DAN we well as decreases in DMN3 and DMN4. Working memory elicited the strongest positive effects in FPN and negative ones in DMN4. Changes in CMRGlu for eyes open and finger tapping paradigms are shown for comparison only as these are identical to figure 2 since only fPET data was available. For a detailed description see figure 2.
